# A duplicated copy of the meiotic gene *ZIP4* preserves up to 50% pollen viability and grain number in polyploid wheat

**DOI:** 10.1101/2021.01.22.427771

**Authors:** Abdul Kader Alabdullah, Graham Moore, Azahara C. Martín

## Abstract

- Although most flowering plants are polyploid, little is known of how the meiotic process evolved to stabilise and preserve polyploid fertility. On wheat polyploidisation, the major meiotic gene *ZIP4* on chromosome 3B duplicated onto 5B and subsequently diverged. This 5B meiotic gene copy (*TaZIP4-B2*) was recently shown to promote homologous pairing, synapsis and crossover, and suppress homoeologous crossover. We therefore suspected that these stabilising effects on meiosis could be important for the preservation of wheat polyploid fertility.
- A CRISPR *Tazip4-B2* mutant was exploited to assess the contribution of the 5B duplicated *ZIP4* copy in maintaining pollen viability and grain setting.
- Analysis demonstrated abnormalities in 56% of meiocytes in the *Tazip4-B2* mutant, with micronuclei in 50% of tetrads, reduced size in 48% of pollen grains and a near 50% reduction in grain number. Further studies showed that most of the reduced grain number resulted from pollination with less viable pollen, suggesting that the stabilising effect of *TaZIP4-B2* on meiosis has a greater consequence in subsequent male, rather than female gametogenesis.
- These studies reveal the extraordinary value of the wheat chromosome 5B *TaZIP4-B2* duplication to agriculture and human nutrition. Future studies should assess whether different *TaZIP4-B2* alleles exhibit variable effects on meiotic stabilisation and/or resistance to temperature change.

## Introduction

Polyploidy occurs in a wide range of species, including fish, flatworms, shrimp, amphibians, flowering plants, wine and brewing yeast (Comai 2005; Otto 2007; Pelé *et al*. 2018; Feliner *et al*. 2020). The molecular mechanisms responsible for meiotic polyploidisation and diploid behaviour are important for ensuring correct chromosome segregation of multiple related chromosomes, production of balanced gametes and hence preservation of fertility. It is surprising that these mechanisms have not been more widely investigated, given their potentially enormous value to mankind (Feliner *et al*. 2020).

Plant polyploidisation is often associated with extensive chromosomal rearrangements and changes in gene content and expression (Osborn *et al*., 2003; Adams & Wendel, 2005; Pelé *et al*., 2018; Mason & Wendel, 2020). Yet analysis of the recently sequenced hexaploid wheat (*Triticum aestivum* L.) genome and wheat RNA seq datasets from over 1000 tissues (including meiocytes), did not reveal extensive gene loss or changes in expression between related (homoeologous) chromosomes following polyploidisation (Ramírez-González *et al*., 2018). Even meiotic genes do not appear to have suffered gene loss, exhibiting mostly balanced expression between copies on related chromosomes (homoeologues) (Alabdullah *et al*., 2018). Thus, hexaploid wheat appears to have suffered less extensive rearrangement, gene loss or altered expression compared to other polyploids. This suggests a more rapid and simple adaption occurring on polyploidisation (tetraploid and hexaploid wheat) ensuring genome stability and fertility. High wheat fertility is important since it is consumed by over 4.5 billion people on the planet, of whom 2.5 billion people are dependent on it (Food and Agriculture Organization of the United Nations, 2017).

It was previously accepted that a locus arising on chromosome 5B during wheat polyploidisation, was responsible for stabilising the wheat genome during meiosis, hence maintaining fertility. This was based on earlier cytogenetic studies of hexaploid wheat lines lacking the whole of chromosome 5B, which when crossed with wild relatives such as rye or *Aegilops variabilis*, exhibited homoeologous crossover between wheat and wild relative chromosomes at metaphase I in the resulting hybrids (Riley & Chapman, 1958; Sears & Okamato, 1958). Deletion of the whole 5B chromosome resulted in the loss of multiple meiotic genes but it was unclear at the time which and how many of these genes needed to be lost to produce the phenotype. However, it was recognised from these early studies that suppression of homoeologous crossover was important for stabilising the wheat genome and maintaining its fertility. In 1971, a study coined the term ‘pairing homoeologous’ (*Ph1*) for this ‘critical locus’ on 5B, responsible for suppressing the homoeologous crossover observed in wheat-wild relative hybrids (Wall *et al*., 1971). Loss of *Ph1* (or of the whole 5B chromosome, as in these studies) allowed homoeologous crossover to take place. The term ‘pairing’ was used synonymously with crossover observed at metaphase I at this time. ‘*Ph1’* became the accepted term to describe the locus responsible for the homoeologous crossover suppression phenotype.

Sears (1977) identified a mutant (named *ph1b*) carrying a deletion of part of chromosome 5B (now known to be 59.3Mb in size, encompassing some 1187 genes (Martín *et al*., 2018)). When the Sears *ph1b* mutant is crossed with wild relatives to form hybrids, crossover between homoeologues is subsequently observed during meiosis in these hybrids. Exploitation of such mutants in crosses with wild relatives has allowed the transfer of traits from wild relatives into wheat, saving the global economy billions of dollars over the years. Later, Roberts *et al*., (1999) observed that mutants carrying deletions in the long arm of chromosome 5B, could be separated into 2 groups by scoring the meiotic configurations at metaphase I of the mutants themselves. The presence or absence of multivalents at metaphase I did not distinguish these 5B deletion mutants. However, it was observed that univalents, rod bivalents and multivalents were present in over 50% of meiocytes at metaphase I in the Sears *ph1b* mutant and also some of the 5B deletion mutants, while the wild type (WT) wheat and the remaining 5B deletion mutants exhibited mainly bivalents at metaphase I in all their meiocytes. Thus, the presence of meiotic abnormalities in over 50% of meiocytes could separate the 5B deletion mutants into two groups (Roberts *et al*., 1999). The presence of multivalents suggested that the initial alignment of chromosomes (now termed pairing) and intimate pairing (now termed synapsis) of chromosomes was disrupted.

Recently the two phenotypes (suppression of homoeologous crossover (*Ph1*) in wheat-wild relatives, and the presence of meiotic abnormalities in 50% of meiocytes in the mutant itself) have been defined using a series of 5B deletions to a 0.5Mb region of chromosome 5B containing a copy of the major meiotic gene *ZIP4* (*TaZIP4-B2*) (Griffiths *et al*., 2006; Al-Kaff *et al*., 2008; Rey *et al*., 2017; Martín *et al*., 2018; Rey *et al*., 2018a). Genome analysis revealed that hexaploid wheat possessed a further three *ZIP4* genes on group 3 chromosomes (*ZIP4 3A* (*TaZIP4-A1*), *ZIP4 3B* (*TaZIP4-B1*) and *ZIP4 3D* (*TaZIP4-D1*)). Analysis by the International Wheat Genome Sequencing Consortium (2018) confirmed that on wheat polyploidisation, *TaZIP4-B2* was derived from *TaZIP4-B1* through a trans-duplication event.

ZIP4 is a protein containing tetratricopeptide repeats (TPRs). Proteins with such tandem TPRs can form folds which assemble protein complexes (Blatch & Lassle, 1999; D’Andrea & Regan, 2003). In *Sordaria* (and budding yeast), ZIP4 is necessary for pairing, synapsis and homologous crossover (Tsubouchi *et al*., 2006; Dubois *et al*., 2019; Pyatnitskaya *et al*., 2019), whereas, in Arabidopsis and rice, *ZIP4* has previously only been reported necessary for homologous crossover (Chelysheva *et al*., 2007; Shen *et al*., 2012). In wheat however, *TaZIP4-B2* promotes homologous pairing, synapsis and crossover (since in the wheat CRISPR *Tazip4-B2* mutant, 50% of meiocytes exhibit meiotic abnormalities, the phenotype reported by Roberts *et al*., (1999) (Rey *et al*., 2018a)). Suppression of homoeologous crossover by *ZIP4* has not previously been reported in any species, however when the CRISPR *Tazip4-B2* deletion mutant is crossed with a wild relative to form a hybrid, homoeologous crossover takes place in the hybrid (Rey *et al*., 2018a), implying a role for *TaZIP4-B2* in homoeologous crossover suppression. Thus, the duplication of *ZIP4* on wheat polyploidisation led to an adaption during meiosis I, preventing meiotic disruption by promoting homologous pairing, synapsis and crossover, and suppressing homoeologous crossover.

Following meiosis I, the reductional division during wheat male meiosis (meiosis II), leads to the formation of tetrads each containing four microspores, which degenerate to release individual uninucleate microspores. An asymmetric mitotic division takes place in each microspore to produce a large vegetative cell and a small generative cell. Subsequently, the small generative cell undergoes a second mitotic division to form a mature trinucleate pollen grain, with one vegetative nucleus and two generative nuclei or sperm cells. Reductional division in wheat female meiosis results in a T-shaped tetrad, containing 4 megaspores. Only one of the megaspores develops into an embryo sac, with the remaining 3 megaspores degenerating. Hence, whilst all four products of meiosis survive on the male side, only one survives on the female side. Meiosis is an essential process for the formation of gametes. Thus, meiotic abnormalities or genetic disruptions are likely to result in reduced fertility. Meiotic abnormalities on the male side may be associated with variable sized and/or inviable pollen grains, and on the female side, with a partial reduction in grain number or complete sterility (Pagliarini, 2000; Sheidai *et al*., 2009; 2010; Dewitte *et al*., 2010; Kumar *et al*., 2010; Jiang *et al*., 2011; Singhal & Kaur, 2011; Kaur & Singhal, 2019).

In the polyploid literature, it is often stated that meiotic adaptation is important for polyploid fertility. However, it has not previously been possible to determine the effect of an actual meiotic adaptation. The availability of a CRISPR deletion mutant for the duplicated *TaZIP4-B2* copy allows us to assess the effect of this meiotic adaptation on the correct segregation of chromosomes, effective production of balanced gametes, and hence preservation of pollen viability and grain number in this major global crop. As part of this assessment, a new pollen profiling method has been developed and exploited to compare pollen profiles of different mutants in hexaploid (and tetraploid) wheat.

## Materials and Methods

### Plant material

Three different *Tazip4-B2* mutants were used in this study: 1) *ph1b*: a hexaploid wheat *T. aestivum* cv. Chinese Spring mutant with a gamma radiation induced 59.3 Mb deletion in the long arm of chromosome 5B, including the *TaZIP4-B2* gene copy (Sears, 1977; Griffiths *et al*., 2006); 2) CRISPR *Tazip4-B2*: a hexaploid wheat *T. aestivum* cv. Fielder mutant with a CRISPR-induced 114 bp deletion in exon 1 of *TaZIP4-B2* leading to the deletion of 38 amino acids (A^104^ to E^141^) from the TaZIP4-B2 protein (Rey *et al*., 2018a); 3) *ph1c*: a tetraploid wheat *T. turgidum* subsp. *Durum* cv. Senatore Cappelli mutant carrying a large deletion in the long arm of chromosome 5B, including the *TaZIP4-B2* gene (Giorgi, 1983; Jampates & Dvorak, 1986; Roberts *et al*., 1999).

### Pollen profiling

Plants were grown in a controlled environment room (CER) at 20 °C (day) and 15 °C (night), with a 16-hr photoperiod and 70% humidity. Ten plants were grown for each genotype. The first spike per plant was labelled for collection of anthers. Mature yellow anthers were collected just before shedding pollen, from five main florets at the middle portion of the spike. Each of three anthers from the same floret were placed in an Eppendorf containing 0.5 ml of 70% ethanol and stored at 4°C for later pollen counts and size measurements. Pollen grains were released from anthers by sonication using Soniprep 150 Plus (MSE, Heathfield, East Sussex, UK) at amplitude 5 for 30 seconds. The sonicated pollen samples were filtered through 200 µm sieves using 100 ml Coulter Isoton II diluent (Beckman Coulter) to eliminate anther debris. Size and number of filtered pollen grains were measured using a Coulter counter (Multisizer 4e, Beckman Coulter Inc.), fitted with a 200 µm aperture tube, with Isoton II diluent (using the following settings: Control mode: volumetric; Analytic volume: 2000 µl; Electrolyte volume: 100 mL; Size bins = 400 from 4 µm to 120 µm; Current: 1600 µA; Stirring speed: 20 CW). For each sample, the measured pollen number distribution over size bins was exported into a csv file, then an R script (Text S1) used to extract and calculate plot differential pollen size distribution and pollen number per anther from the raw data files for each genotype.

### Pollen viability

Pollen viability was assessed using Alexander stain (Alexander, 1969). Briefly, Alexander stain was prepared according to Alexander (1969). Fresh wheat pollen grains from three anthers were shed on a droplet of Alexander stain placed on a microscopic slide and covered with a coverslip for microscopic observation. Images were taken from several random microscope field views to be scored. Magenta-coloured pollen was considered viable, whereas blue-green pollen was considered to be non-viable or sterile. Three biological replicates with >1000 pollen grains each were analysed for each genotype.

### Grain number per spike assessment

The effect of *Tazip4-B2* mutants on grain number per spike (or grain setting) was assessed under CER and glasshouse conditions. In the CER, plants from each of the four genotypes CRISPR *Tazip4-B2, ph1b* and their corresponding WTs, were grown at 20 °C (day) and 15 °C (night) with a 16-hr photoperiod and 70% humidity. In the glasshouse, 11-15 plants from each of the CRISPR *Tazip4-B2* and *ph1b* mutants, with their corresponding WTs, were grown at 22 °C (day) and 17 °C (night) with an 8-hr photoperiod and 70% humidity. In both experiments, the first three spikes from each plant were tagged, bagged at the heading stage, harvested when fully dried, and threshed separately after counting spikelet number. The number of grains per spike was then measured using the MARVIN grain analyser (GTA Sensorik GmbH, Neubrandenburg, Germany). Grain number per spike ((actual grain number per spike/expected grain number per spike)^*^100) was normalised in order to eliminate the effect of different number of spikelets per spike on grain number. Expected grain number was calculated by multiplying number of spikelets by three, considering that each spikelet has three main fertile florets.

### Female sterility assessment

Female sterility was assessed through the emasculation/pollination method, using the CRISPR *Tazip4-B2* mutant as both pollen donor and recipient. Three treatments were involved in this experiment: 1) Using CRISPR *Tazip4-B2* mutant as the ‘pollen recipient’, where the mutant was pollinated with pollen from the WT *T. aestivum* cv. Fielder (pollen donor), 2) Using CRISPR *Tazip4*-*B2* mutant as a ‘pollen donor’, where WT plants were pollinated with CRISPR *Tazip4-B2* pollen; 3) Emasculating WT plants and pollinating them with WT pollen, providing a control for measurement of emasculation and pollination procedure efficiency. For each pollination experiment, at least twelve spikes were emasculated at the heading stage when the spike had fully emerged from the flag leaf and anthers were still green with a tight stigma. Spikelets located at the tip and base of the spike and florets in the centre of each spikelet were removed before emasculation, as they are usually asynchronous to the rest of the spike and frequently sterile. Emasculated spikes were covered with crossing bags to avoid dehydration and any unwanted cross-pollination. When the emasculated floret stigma was receptive and mature (usually 2-3 days after emasculation), pollination was performed using fresh pollen grains collected from fully mature anthers just at opening stage. All emasculations and pollinations were undertaken at the same time of day (in the morning) to avoid any possible circadian effects on stigma and pollen fertility. Grain number was then counted for each spike and normalised by dividing the number of grains by the number of pollenated florets per spike.

### Seed germination rate

Germination rates of seeds resulting from the emasculation/pollination experiment were evaluated to assess the paternal and maternal effect of the *Tazip4-B2* mutant on embryo development after fertilization. Before gemination, seeds were disinfected by soaking in 5% Sodium Hypochlorite for 5 minutes and then washed with distilled water. Seeds were placed in Petri dishes (9 cm diameter) containing two layers of filter paper wetted continuously with distilled water. Each Petri dish represented a replicate containing 15 seeds originating from the same spike. Five replicates were used for each treatment. Petri dishes were wrapped with aluminium foil and placed in a growth chamber at 22°C. The seeds were considered to have germinated after radicle emergence. Germination period was 10 days. The germination percentage was calculated as (number of seeds germinated /total seeds) × 100.

### Meiotic analysis

Anthers were collected at the desired meiotic stage as previously described (Martín *et al*., 2017) and fixed in freshly prepared 100% ethanol/glacial acetic acid 3:1 (v/v). The material was transferred to fresh fixative after 1–2 hr and stored at 4 °C until needed for Feulgen or FISH (fluorescence *in situ* hybridisation) studies. Cytological analysis of Pollen Mother Cells using the Feulgen technique was performed as previously described (Sharma & Sharma, 2014). Anthers fixed at the tetrad stage were used for FISH analysis. Preparations were made as described by Rey *et al*., (2018b). The repetitive sequence 4P6 (Zhang *et al*., 2004) was amplified by PCR as previously described (Rey *et al*., 2018b) and labelled using the DIG-nick translation mix (Sigma, St. Louis, MO, USA) according to the manufacturer’s instructions. The repetitive probe pTa71, containing 1 unit of 18S-5.8S-26S rDNA (8.9 kb) from *T. aestivum* (Gerlach & Bedbrook, 1979) was labelled using the Biotin-nick translation mix (Sigma). Digoxigenin-labelled probes were detected with anti-digoxigenin-fluorescein Fab fragments (Sigma) and Biotin-labelled probes were detected with Streptavidin-Cy5 (Thermo Fisher Scientific, Waltham, Massachusetts, USA). FISH was performed as described previously (Rey *et al*., 2018b).

### Image processing

Pollen grains and Pollen Mother Cells stained by the Feulgen technique were imaged using a LEICA DM2000 microscope (Leica Microsystems, http://www.leica-microsystems.com/), equipped with a Leica DFC450 camera and controlled by LAS v4.4 system software (Leica Biosystems, Wetzlar, Germany). Tetrads labelled by FISH were imaged using a Leica DM5500B microscope equipped with a Hamamatsu ORCA-FLASH4.0 camera and controlled by Leica LAS X software v2.0. Z-stacks were processed using the 561 deconvolution module of the Leica LAS X Software package. Images were processed using Adobe Photoshop CS5 (Adobe Systems Incorporated, US) extended version 12.0 × 64.

### TaZIP4 proteins sequence analyses

The DNA, CDs and protein sequences of the four *TaZIP4* homologues were retrieved from the *Ensembl Plants* database for *Triticum aestivum* (IWGSC v1.1 gene annotation; International Wheat Genome Sequencing Consortium, 2018). Multiple sequence alignments of coding sequences (CDs) and protein sequences of *TaZIP4-A1* (TraesCS3A02G401700.3), *TaZIP4-B1* (TraesCS3B02G434600.2), *TaZIP4-B2* (TraesCS5B02G255100.1), *TaZIP4-D1* (TraesCS3D02G396500.2) and the mutant CRISPR Tazip4-B2 (Rey *et al*., 2018a) were performed using the Clustal X programme (version 2; Higgins & Sharp, 1988; Larkin *et al*., 2007). Functional domain prediction in the protein sequences was performed using the online InterPro programme (version 82.0; Mitchell *et al*., 2019). Tetratricopeptide Repeats (TPRs) in the TaZIP4 proteins were predicted using the online TPRpred program (version 11.0; Magis *et al*., 2014; Zimmermann *et al*., 2018). Prediction of coiled coil domains in the TaZIP4 proteins was performed using the MARCOIL programme (Delorenzi & Speed, 2002; Zimmermann *et al*., 2018).

## Results

### Divergence and the CRISPR deletion occur within the TaZIP4-B2 TPR domain

The InterProScan (Mitchell *et al*., 2019) and PFAM programmes identified a single highly conserved SPO22 domain (PF08631) within the EBI database ZIP4s. This SPO22 domain was composed of tetratricopeptide repeats (TPRs) of 34 amino acids. A second SPO22 domain of low significance was observed in tandem with the highly conserved SPO22 domain in many ZIP4s. Only PFAM classified this second SPO22 domain as being significant for a limited number of these ZIP4s. ZIP4 function is dependent on these SPO22 Tpr-containing domains, due to their involvement in assembling protein complexes (Blatch & Lassle, 1999; D’Andrea & Regan, 2003). The annotation programmes enabled us to assess whether the divergence of *TaZIP4-B2* from its chromosome group 3 homoeologues (*TaZIP4-A1, TaZIP4-B1* and *TaZIP4-D1*) occurred within the SPO22 domain. Similarly, we used the annotation programmes to determine the site of the in-frame 38 amino acid CRISPR deletion of *Tazip4-B2* (Rey *et al*., 2018a) relative to the SPO22 domain. Multiple sequence alignments showed that TaZIP4-B2 was quite divergent from the other group 3 homoeologues (Fig. 1a). The percentage of identity between TaZIP4-B2 and the other homoeologues did not exceed 85.8% in coding sequences (CDs) and 92.2% in protein sequences, whereas the inter-identity of TaZIP4-A1, TaZIP4-B1 and TaZIP4-D1 ranged from 94.9% - 96.3% for CDs sequences and from 96.8% - 97.5% for protein sequences (Table S1). The InterProScan and PFAM programmes identified the highly conserved SPO22 domain within all the wheat ZIP4s, with PFAM identifying a second SPO22 domain in tandem (Fig. 1b). TPRpred (Zimmermann *et al*., 2018) identified 12 TPRs within wheat ZIP4-B1 (Fig. 1c), showing that up to half of total ZIP4-B1 protein consisted of TPRs. However, the region of TaZIP4-B2 corresponding to the 3^rd^ TPR of ZIP4-B1 within the highly conserved SPO22 domain, was no longer identified as a TPR by TPRpred. Thus, within wheat TaZIP4-B2, only 11 TPRs were identified. The 2^nd^ and 4^th^ TPRs of TaZIP4-B2 and TaZIP4-B1 also exhibited some divergence with respect to each other. As a result of this divergence, the MARCOIL programme (Zimmermann *et al*., 2018; Delorenzi & Speed, 2002) suggested an altered conformation within the conserved SPO22 domain of TaZIP4-B2 compared to the domains of TaZIP4-B1 and other ZIP4 homoeologues (Fig. 1d). Thus, duplication of *TaZIP4-B2* from *TaZIP4-B1* led to TPR divergence (especially the 3^rd^ TPR (Table S1)), giving rise to associated changes in protein conformation. This predicts that *TaZIP4-B2* function may be altered with respect to that of its chromosome group 3 homoeologues. The 38 amino acid in-frame CRISPR deletion (within the CRISPR *Tazip4-B2*) covered the 1^st^ TPR, indicating that the deletion did indeed affect the SPO22 domain and correlated with complete loss of the *TaZIP4-B2* phenotype (Rey *et al*., 2018a).

**Fig 1.**
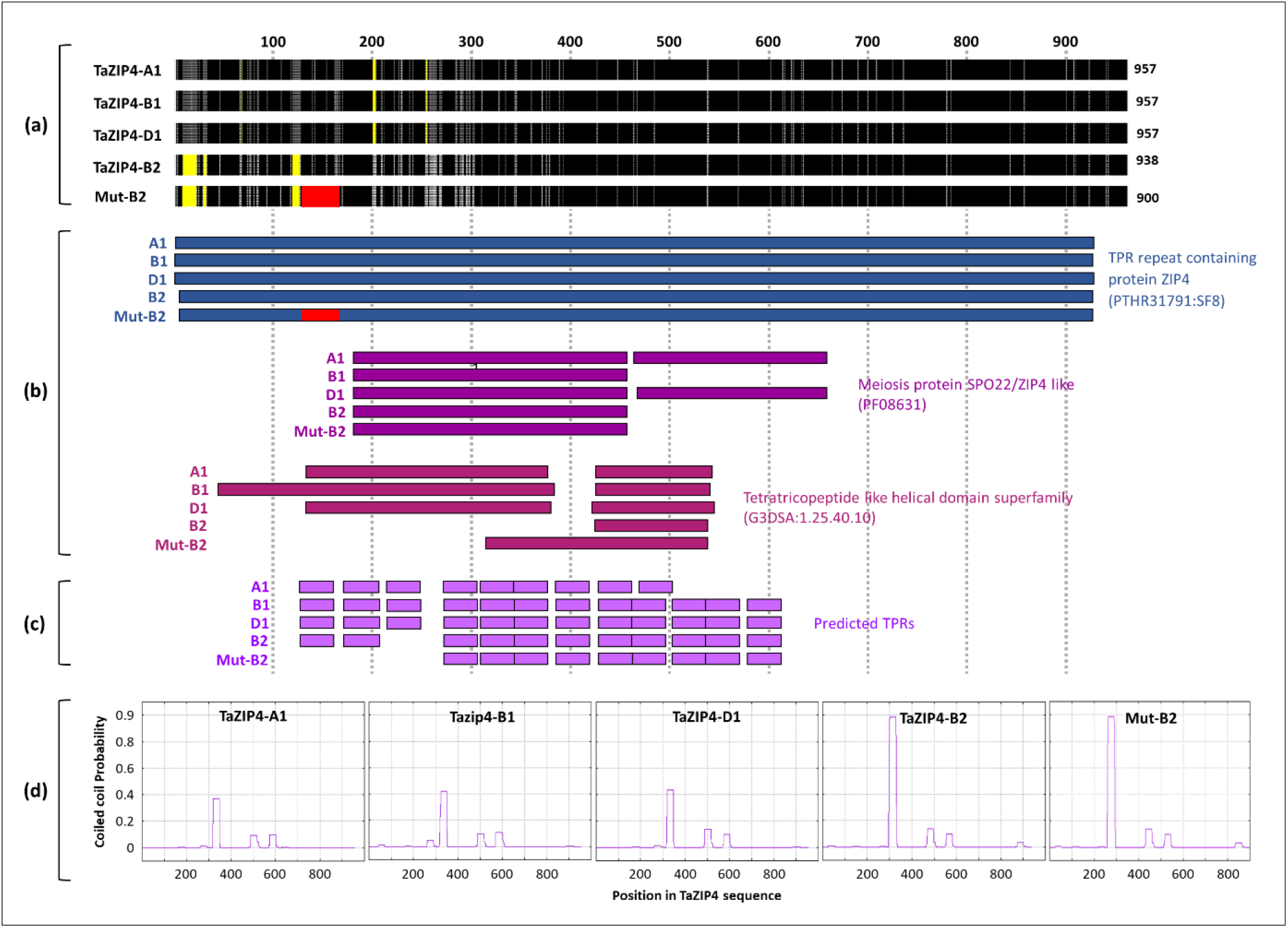
Comparison of the TaZIP4 homoeologous proteins. **(a)** Multiple amino acid sequence alignment of the TaZIP4 homoeologous proteins using ClustalX software (version 2.0; Higgins & Sharp, 1988; Larkin *et al*., 2007). Regions with identical amino acid sequences across the four proteins are in black. Grey colour refers to the sequences with similar amino acid properties and light grey refers to sequences with different amino acid properties. Yellow regions indicate gaps in the sequence alignment. Mut-B2 refers to *Tazip4-B2* in the CRISPR mutant. Red region shows the 38-amino acids segment that is deleted from the protein of CRISPR *Tazip4-B2* mutant. **(b)** Predicted functional domains in the TaZIP4 proteins using the online InterPro software (version 82.0; Mitchell *et al*., 2019). **(c)** The predicted Tetratricopeptide Repeats (TPRs) in the TaZIP4 proteins using the online TPRpred program (version 11.0; Magis *et al*., 2014; Zimmermann *et al*., 2018). **(d)** The predicted coiled coil domains in the TaZIP4 proteins using the online MARCOIL programme (Delorenzi & Speed, 2002; Zimmermann *et al*., 2018).

### Effect of the *TaZIP4-B2* deletion on meiotic and tetrad stages

Average meiotic scores from *Tazip4-B2* mutants at metaphase I were reported previously (Martín *et al*., 2014; Rey *et al*., 2018a). However, the present study required meiotic scores from individual meiocytes from *Tazip4-B2* mutants at metaphase I, in order to relate meiotic abnormalities observed during metaphase I with those observed at subsequent stages. Both CRISPR *Tazip4-B2* (Rey *et al*., 2018a) and *ph1b* (Sears, 1977) mutants were exploited. The *ph1b* mutant carries a 59.3Mb deletion encompassing some 1187 genes, including *TaZIP4-B2* (Martín *et al*., 2018). Meiotic scores from individual meiocytes at metaphase I from CRISPR *Tazip4-B2* (Rey *et al*., 2018a), *ph1b* (Martín *et al*., 2014) and their respective wild type plants are provided in Table 1, Table S2 and visualised in Fig. 2. Examples of meiotic configurations of the CRISPR *Tazip4-B2* mutant and wild type (WT Fielder) plants at metaphase I are provided in Fig. 3a, b. More than half of the scored meiocytes in the CRISPR *Tazip4-B2* and *ph1b* mutants had meiotic abnormalities (Fig. 2). Overall, univalents and/or multivalents were observed in 56% of both the CRISPR *Tazip4-B2* and *ph1b* mutant meiocytes. Univalents were present in 49% and 43% of meiocytes at metaphase I (average per meiocyte 1.16 and 0.8) for the CRISPR *Tazip4-B2* and *ph1b* mutants respectively. Multivalents were present in 32% and 43% (average per meiocyte 0.39 and 0.53) of the CRISPR *Tazip4-B2* and *ph1b* mutant meiocytes respectively. The slight excess of multivalents and lack of univalents in the *ph1b* mutant compared to the CRISPR *Tazip4-B2* mutant may be simply due to accumulation of the extensive rearrangements observed and reported in this mutant (Martín *et al*., 2018), which can form multivalents at metaphase I (Table 1; Table S2). The excess of multivalents in the *ph1b* mutant could also be explained by additional unknown deleted genes within the 59.3Mb 5B deletion, but the issue cannot be resolved by just scoring for the presence or absence of multivalents at metaphase I, as scoring multivalents alone would fail to distinguish between different deletion mutants covering variable lengths of the long arm of 5B (Roberts *et al*., 1999). Thus the deletion of *TaZIP4-B2* leads to nearly half of meiocytes possessing univalents as a result of pairing and crossover failure, and a third of meiocytes possessing multivalents as a result of incorrect pairing and crossover.

**Table 1.** Summary of the meiotic scores for the two *Tazip4-B2* mutants and the corresponding wild types.

| Genotype | Univalents | Multivalents | Rod bivalents | Ring bivalents | Defective meiocytes* (%) | Reference |
| --- | --- | --- | --- | --- | --- | --- |
| Fielder WT** | 0.16±0.07 | 0±0 | 1.37±0.13 | 19.52±0.14 | 6.70 | Rey *et al.*, 2018 |
| CRISPR *Tazip4-B2* | 1.16±0.12 | 0.39±0.05 | 4.93±0.15 | 14.84±0.19 | 56.00 |  |
| Chinese Spring | 0±0 | 0±0 | 1±0.20 | 20±0.20 | 0.00 | Martín *et al.*, 2014 |
| *ph1b* | 0.80±0.19 | 0.53±0.12 | 4.73±0.26 | 14.83±0.33 | 56.60 |  |

\* Meiocytes with meiotic aberrations (univalents and multivalents) thus have incorrect chromosomes paring.
\*\* This is a CRISPR transgenic Fielder without *TaZIP4-B2* knockout.

**Fig 2.**
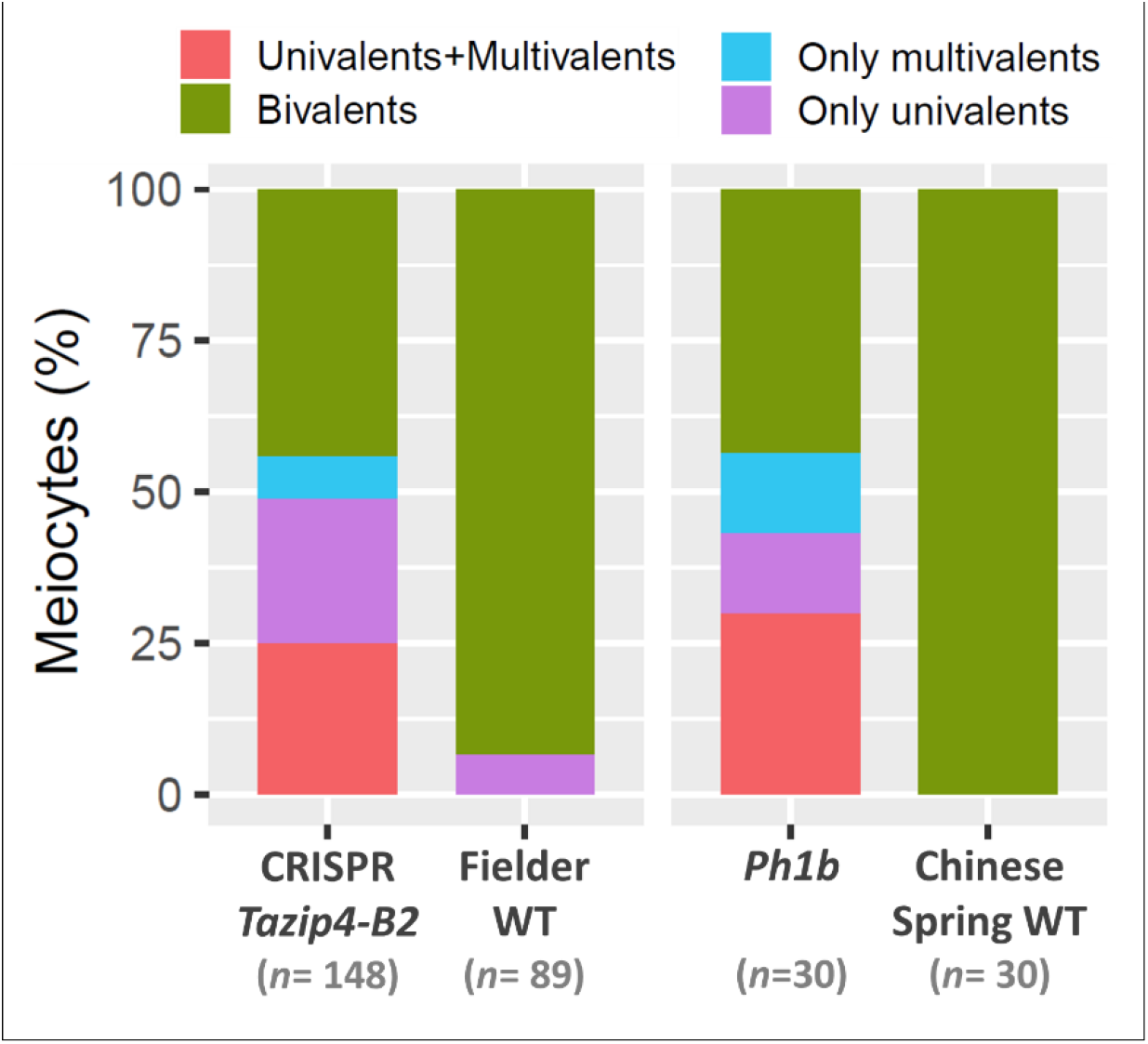
The percentage of meiocytes with meiotic abnormalities from the CRISPR *Tazip4-B2* and *ph1b* mutants, and their wild types. The data used to produce this figure is taken from Martín *et al*., 2014 and Rey *et al*., 2018. *n* refers to the number of scored meiocytes.

**Fig 3.**
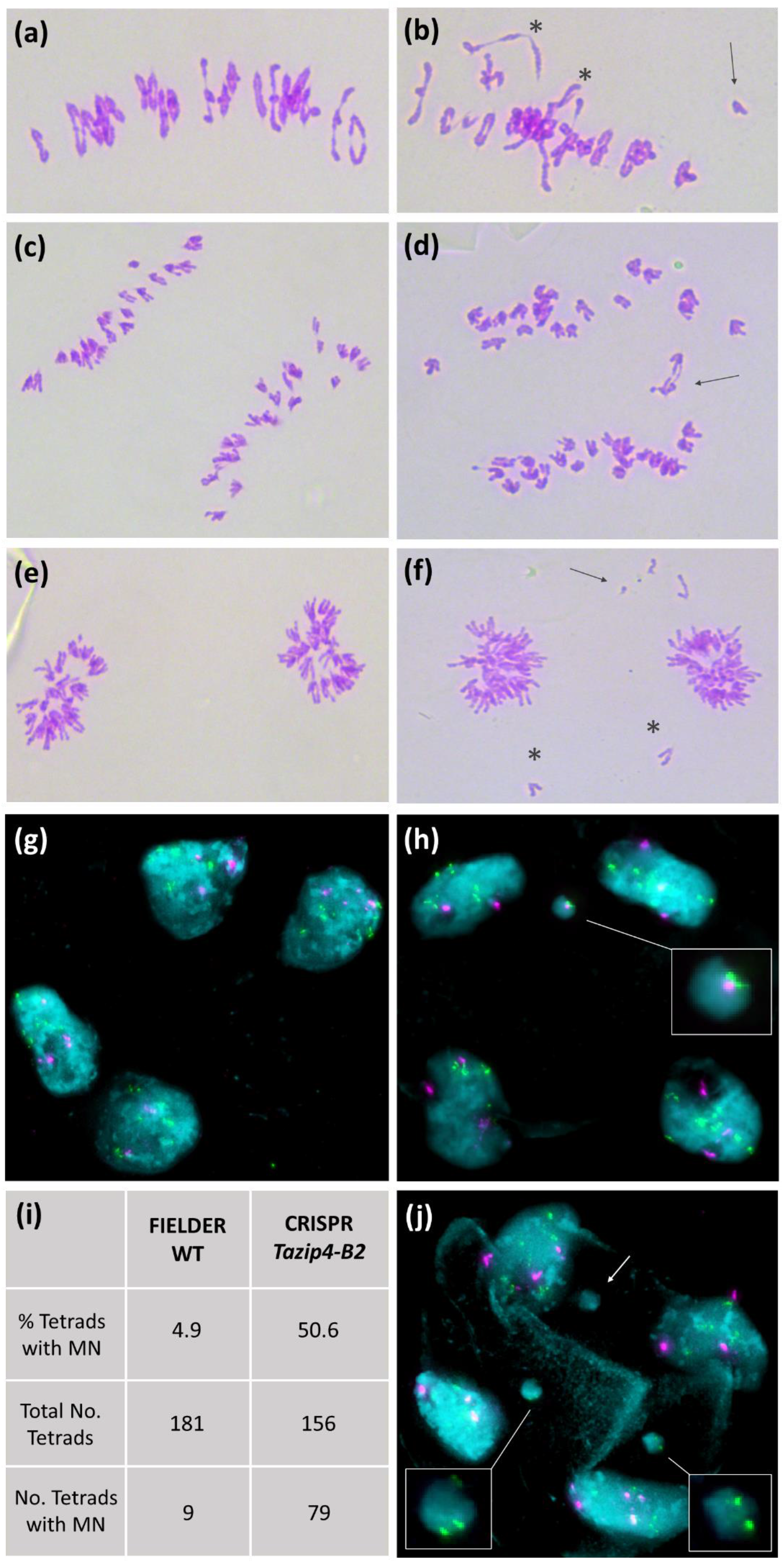
Meiosis in wild type (WT) Fielder (a,c,e,g) and the CRISPR *Tazip4-B2* Fielder mutant (b,d,f,h,j). (**a**) Metaphase I in WT Fielder showing 19 ring bivalents and 3 rod bivalents. (**b**) Metaphase I in CRISPR *Tazip4-B2* mutant with the presence of multivalents (asterisk) and univalent (arrow). (**c**) Anaphase I in WT displaying equal separation of homologous chromosomes to both poles. (**d**) Anaphase I in CRISPR *Tazip4-B2* mutant showing lagging chromosomes. (**e**) Late anaphase I in WT. (**f**) Late anaphase I in CRISPR *Tazip4-B2* mutant showing some chromosome fragments in the periphery of equatorial plate (arrow) and chromosome mis-division in the equatorial plate (asterisk) which will not be included in any of the diads. (**g, h**) Tetrads shown in cyan, with repetitive probe 4P6 (in green) and pTa71 (in magenta). (**g**) Tetrad from WT showing 4 normal microspores. (**h**) Tetrad in CRISPR *Tazip4-B2* mutant showing 1 micronuclei (MN) displaying 4P6 and pTa71 signals. (**j**) Tetrad in CRISPR *Tazip4-B2* mutant showing 3 micronuclei, 2 of them presenting 4P6 signals. (**i**) Close to 5% of the tetrads show MN in WT, while 50% of the tetrads in the CRISPR *Tazip4-B2* mutant possess 1, 2 or 3 MN.

Meiotic aberrations at metaphase I (univalents and multivalents) can lead to imbalanced chromosomal segregation at anaphase I, with subsequent disruption to the post-meiotic process. Therefore, the stages following metaphase I were studied in both the WT Fielder and the CRISPR *Tazip4-B2* mutant. In WT Fielder, homologous chromosomes (homologues) appear connected to each other by one or mostly several crossovers (Fig. 3a), with only an occasional univalent being present during metaphase I. Each homologue separates to a different pole of the nucleus during anaphase I, resulting in equal separation of homologues (Fig. 3c, e). After the second meiotic division, tetrads with four balanced gametes each are formed (Fig. 3g). In the CRISPR *Tazip4-B2* mutant, univalents, multivalents and a global reduction in the number of crossovers were observed at metaphase I (Fig. 3b), as previously reported (Rey *et al*., 2018a). Although unbalanced segregation of chromosomes would be expected during anaphase I as a consequence of disrupted crossover distribution, disruptions observed were greater than expected, with regular presence of lagging chromosomes, split sister chromatids and chromosome fragmentation (Fig. 3d, f). The high number of micronuclei (MN) observed in tetrads, the final product of meiosis, was the most surprising result (Fig. 3h, j). MN are formed as a consequence of laggard chromosomes or fragments from mis-division that have not been included in telophase I nuclei and are maintained during the second meiotic division (Morrison, 1953). It is not unusual to find an occasional MN in wheat. Indeed, some were found in the WT Fielder analysed in this study (less than 5% of tetrads), probably due to an occasional univalent observed at metaphase I. However, in the CRISPR *Tazip4-B2* mutant, it was striking that more than 50% of tetrads showed at least one MN (Fig. 3i); one, two and less frequently three MN per tetrad were detected. Fluorescence *in situ* hybridisation (FISH) was performed on tetrads from both the WT Fielder and CRISPR *Tazip4-B2* mutant, using the repetitive probes 4P6 (Zhang et al., 2004) and pTa71 (Gerlach and Bedbrook, 1979), in order to assess the level of mis-segregation and to ascertain whether specific chromosomes were involved in MN formation. Probe 4P6 labels seven interstitial sites on D genome metaphase I chromosomes, while pTa71 labels the NOR (Nucleolar Organiser Region) on the 1BS, 6BS and 5DS metaphase I chromosomes. A 4P6 signal was observed in 23.8% MNs, confirming a D genome chromosome origin, and a pTa71 signal in 17.5 %MNs, indicating that some chromosomes were carrying a NOR. This suggests that MN formation did not result from a single specific pair of homologues being univalent at metaphase I, but rather from different pairs of homologues being univalent in individual meiocytes. Morrison (1953) observed that univalents at metaphase I lagged at anaphase I, and then formed MN at the dyad stage. Such MNs were then maintained until the tetrad stage, when they were lost with the separation of the four microspores. Morrison (1953) also observed a direct correlation between numbers of univalents at metaphase I and percentage of tetrads with MN. As such, our observations are consistent with those of Morrison (1953), in that 56% of *Tazip4-B2* mutant meiocytes exhibited abnormalities at metaphase I, while 50% of tetrads subsequently possessed MN.

### Effect of the *Tazip4-B2* deletion on wheat grain number per spike (grain setting)

The presence of MN in 50% tetrads suggested unbalanced microspores, which could also affect grain set. Two experiments (CER and glasshouse) were therefore conducted to assess the effect of deleting *Tazip4-B2* on grain set. In these experiments, grain setting analysis was performed on both the CRISPR *Tazip4-B2* mutant and the *ph1b* hexaploid wheat mutant carrying the 59.3Mb deletion covering *Tazip4-5B*. Spikelet number was recorded, as well as number of grains per spike for the first three spikes from each mutant and their corresponding WTs. The normalized grain number per spike was used to compare genotypes. Both CER and glasshouse experiments confirmed significantly reduced seed set in both *Tazip4-B2* mutants compared to the corresponding WT (*P*<0.01) (Fig. 4a; Table 2). Under CER conditions, the grain number per spike was reduced by 36% in the CRISPR *Tazip4-B2* compared to the WT Fielder, and by 42% in the *ph1b* mutant compared to the Chinese Spring WT. Under glasshouse conditions, the grain number per spike was reduced by 44% in the CRISPR *Tazip4-B2* and 43% in the *ph1b* mutant, compared to their corresponding WTs. There was no significant difference between the CER and glasshouse growth conditions on grain settings for each genotype (Table 2; Table S3). Thus, the CRISPR deletion of *TaZIP4-B2* in hexaploid wheat resulted in 56% of meiocytes exhibiting meiotic abnormalities, 50% of tetrads exhibiting micronuclei, and up to 44% reduction in grain set. Similarly, the *ph1b* mutant also exhibited 56% meiocytes with meiotic abnormalities and up to 43% reduction in grain set.

**Table 2.** Mean Normalized grain number per spike for the two *Tazip4-B2* mutants and their corresponding wild types under CER and glasshouse growth condition. N is the number of biological replicates per genotype. Mean values with standard deviation are shown. Treatments with the same letter are not significantly different.

| Genotype | Controlled Environment Room (CER) |  | Glasshouse |  |
| --- | --- | --- | --- | --- |
|  | N | Normalized grain number per spike | N | Normalized grain number per spike |
| CRISPR *Tazip4-B2* | 5 | 50.2 ± 16.7<sup>bcd</sup> | 15 | 37.7±1.0<sup>de</sup> |
| cv. Fielder WT | 12 | 78.5±12.9<sup>a</sup> | 15 | 67.6±07.9<sup>ab</sup> |
| *ph1b* | 15 | 46.7±1.0<sup>cd</sup> | 11 | 43.8±08.6<sup>cd</sup> |
| cv. Chinese Spring WT | 20 | 80.9±17.3<sup>a</sup> | 15 | 77.3±08.8<sup>a</sup> |

**Fig 4.**
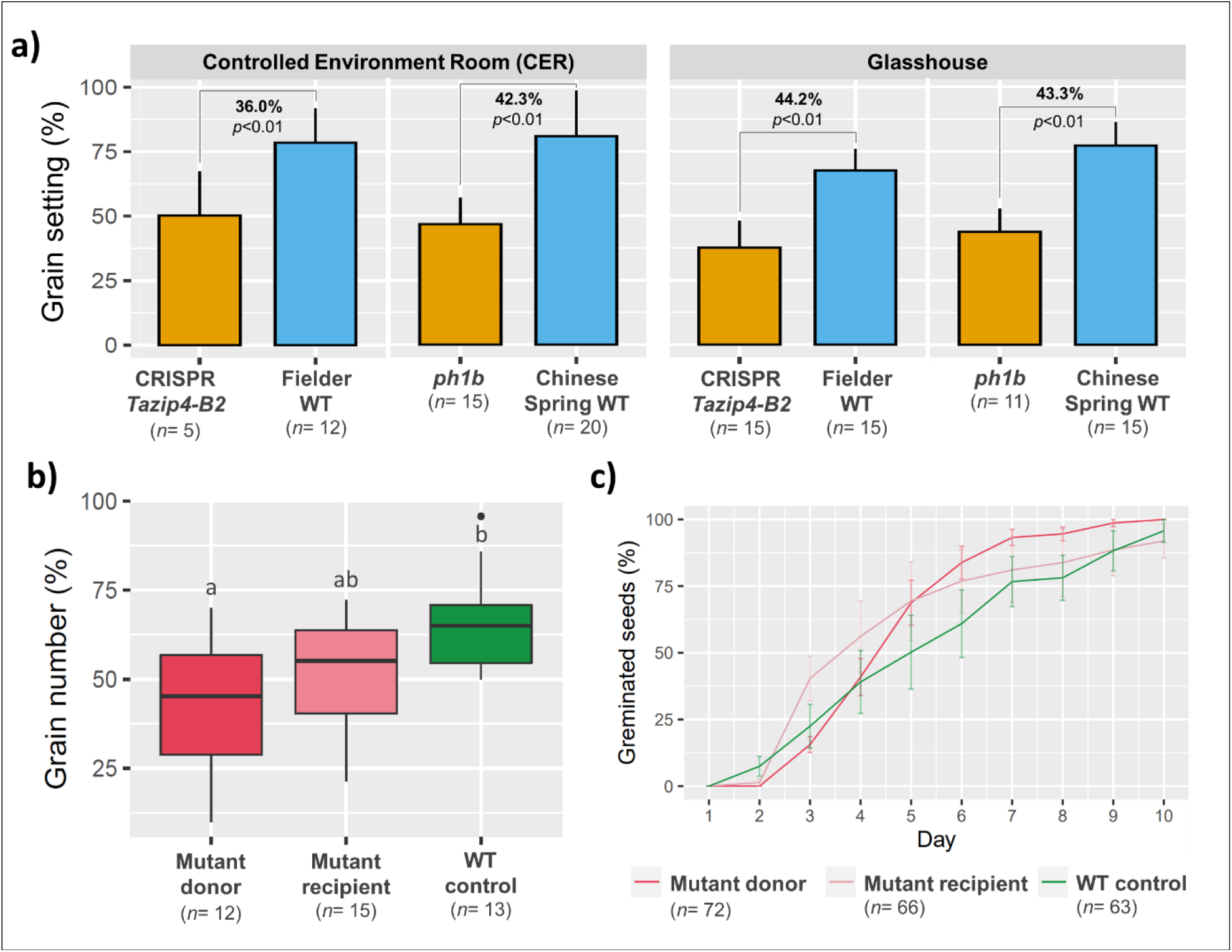
The effect of *TaZIP4-B*2 on grain setting. **a)**. Grain number per spike in the two *Tazip4-B2* mutants and their WT controls under the CER and glasshouse growth conditions. The percentages indicate the difference in grain setting between each mutant and its WT. *n* refers to the number of biological replicates. **b)** The normalised grain number per spike in the three treatments of the emasculation/pollination experiment. Treatments with the same letter are not significantly different. *n* refers to the number of emasculated/pollenated spikes. **c)** Seed germination rates of the seeds resulted from different pollen donor and pollen recipient genotypes. *n* refers to the number of seeds included in the seed germination experiment.

### Pollen contributes to the *Tazip4-B2* effect on grain setting

As previously described, on the female side, only one of the 4 megaspores develops into the embryo sac, with the 3 remaining megaspores degenerating following the tetrad stage. This contrasts with the male side, where all four products of meiosis survive to go through pollen development. It is possible that on the female side, some of the unbalanced megaspores are aborted, so that the near 50% reduction in grain set in the CRISPR *Tazip4-B2* mutant mostly results from pollination with less viable pollen. An emasculation/pollination experiment was therefore conducted, using the CRISPR *Tazip4-B2* mutant and its corresponding WT. The experiment involved pollinating WT plants with WT or *Tazip4-B2* mutant pollen, or the *Tazip4-B2* mutant with WT pollen. Results showed that the lowest percentage of grain number per spike occurred when WT plants were pollinated with CRISPR *Tazip4-B2* mutant pollen (Fig. 4b; Table S4), and that this grain set was significantly lower than that produced by pollinating WT plants with WT pollen (37.8% difference; *P* < 0.01) (Fig. 4b). In contrast, when the *Tazip4-B2* mutant was pollinated with WT pollen, the reduction in grain set was not significantly different to when WT plants were pollinated with WT pollen (Table S4). These results show that most of the reduced grain number in the CRISPR *Tazip4-B2* mutant may be due to its being pollinated with less viable pollen, rather than it all being due to impaired female gametogenesis. Thus, meiotic abnormalities associated with *TaZIP4-B2* deletion may have a greater subsequent effect on male gametogenesis than on female gametogenesis.

The maternal and paternal effects of *TaZIP4-B2* on seed embryo development were assessed by germinating the resulting seeds from each of the above pollination experiments. Germination rates from each of the pollination experiments were not significantly different (Fig. 4c; Table S5). Thus there was no apparent negative effect of the CRISPR *Tazip4-B2* mutation on the germination of seed derived from WT plants pollinated with *Tazip4-B2* mutant pollen, or from*Tazip4-B*2 mutants pollinated with WT pollen.

### A new pollen profiling approach reveals 50% *Tazip4-B2* mutant pollen is small

Meiotic abnormalities in 56% meiocytes lead to mis-segregation of chromosomes and 50% tetrads with micronuclei. The *Tazip4-B2* mutant has up to a 44% reduction in grain number. The emasculation and pollination experiment suggests that most of this effect is the result of reduced pollen viability. We therefore developed a new pollen profiling approach in order to facilitate the study of any effect of the CRISPR *Tazip4-B2* mutant on wheat pollen size and number. The method was validated using pollen samples from five different wheat varieties, namely Cadenza, Fielder and Paragon (hexaploid), Cappelli and Kronos (tetraploid) and one hexaploid wheat landrace (Chinese Spring). Fully mature anthers were collected from the middle portion of the first ear of each plant, just before opening and pollen shedding, and stored in 70% ethanol. The samples could be stored in ethanol for a long period before analysis, without significant effect on pollen measurement accuracy. Pollen profiles of anther samples in 70% ethanol from the same genotype after different storage periods (of up to one month) are shown in Fig. S1. Sonication was used to ensure that all pollen grains were released from anthers, ensuring accurate measurement of pollen number per anther. Pollen size measurements from the six wheat varieties showed that the average pollen size in the hexaploid wheats was 49.0±0.4 µm, (ranging from 48.6±1.2 µm to 49.5±1.1 µm in Chinese Spring and Paragon respectively), while in the tetraploid wheats it was 44.6±0.2 µm (44.8±1.4 µm and 44.4±1.4 µm in Cappelli and Kronos respectively) (Table 3; Table S6), in keeping with previously reported wheat pollen sizes (Cetl, 1960; Saps, 2021)). Pollen profiles of the hexaploid and tetraploid wheat varieties are shown in Fig. 5a.

**Table 3.**
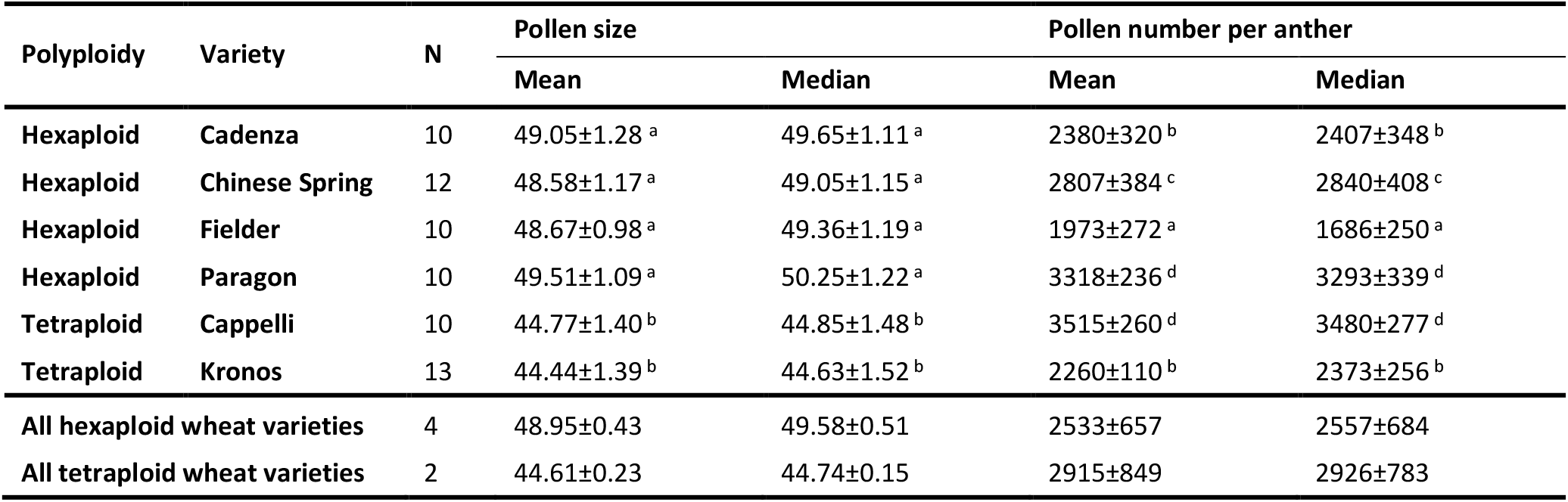
Pollen number and pollen grain size for some hexaploid and tetraploid wheat varieties. Mean and median values with standard deviation are shown. Treatments with the same letter are not significantly different.

**Fig 5.**
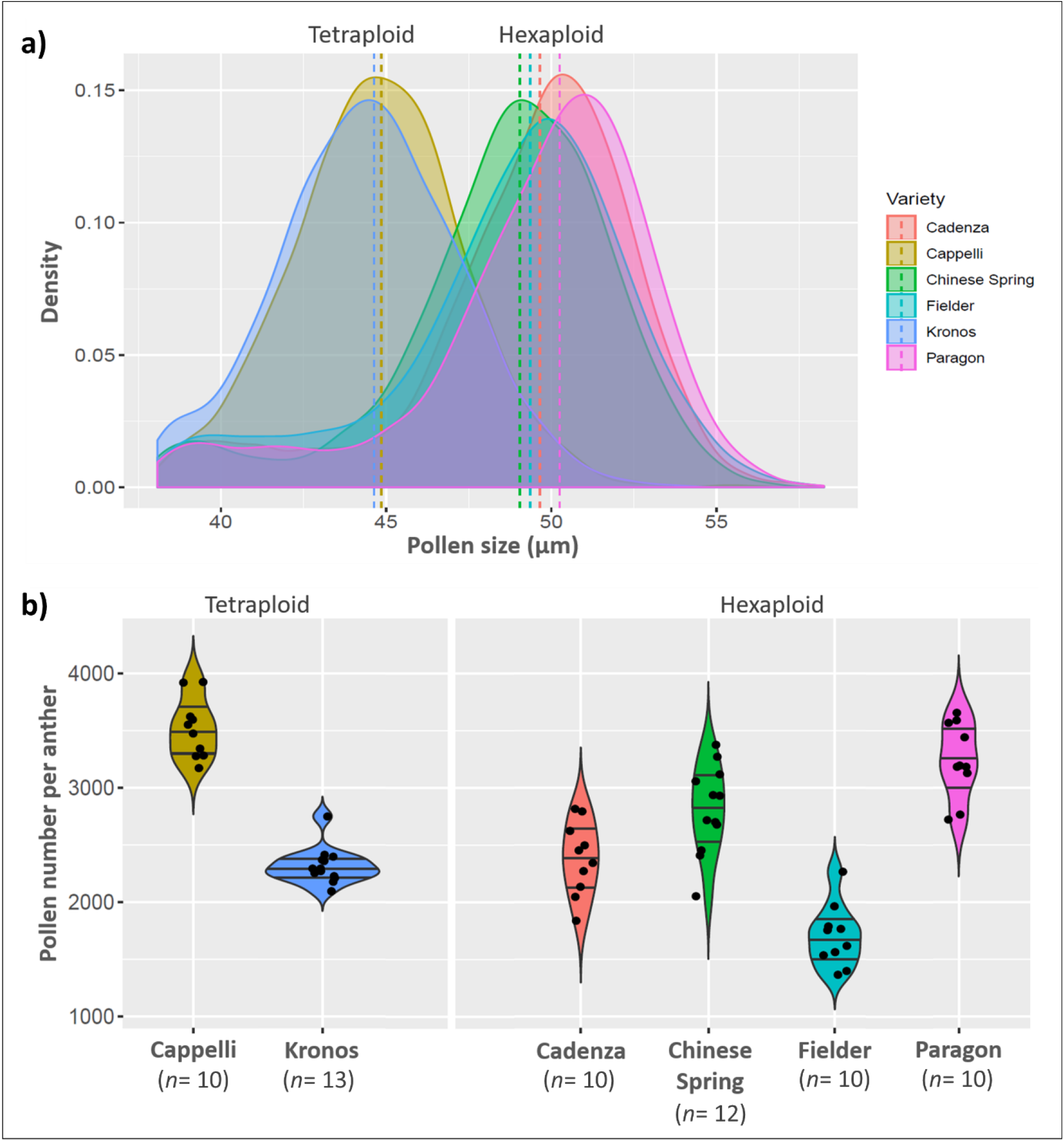
Pollen size and number per anther of some hexaploid and tetraploid wheats. **a)** Density plot of the differential pollen size distribution data collected by coulter counter (Multiziser 4e) for four hexaploid wheats (Chinese Spring, Cadenza, Fielder and Paragon) and two tetraploid wheats (Cappelli and Kronos). Dotted lines indicate the median pollen grain size for each genotype. **b)** Pollen number per anther for the six mentioned hexaploid and tetraploid wheat varieties. *n* is number of plants (biological replicates).

Pollen number per anther varied between different wheat varieties (Fig. 5b). The average number of pollen grains per anther was 2709±614 (ranging from 1973±272 in Fielder to 3515±260 in Cappelli) (Table 3). There was no correlation between number of pollen grains and polyploidy level, as there were no significant differences between hexaploid wheats (Cadenza and Paragon) and tetraploid wheats (Kronos and Cappelli) (Table S6). Nevertheless, pollen numbers were in keeping with those reported in a previous study (De Vries, 1974). The pollen profiling method allowed us to compare pollen grain size distribution and pollen number from three different *Tazip4-B2* mutants with the relevant WT controls. Pollen was collected from full mature anthers (just before opening) for each of the *Tazip4-B2* mutants (CRISPR *Tazip4-B2; ph1b* hexaploid wheat mutant carrying a 59.3Mb chromosome 5B deletion covering *TaZIP4-B2*; *ph1c* tetraploid mutant carrying a large deletion of chromosome 5B covering *TaZIP4-B2*) and their WTs (*T. aestivum* cv. Chinese Spring; *T. turgidum* subsp. *Durum* cv. Senatore Cappelli (Giorgi, 1983); *T. aestivum* cv. Fielder respectively). Ten to twelve biological replicates for each of the six genotypes were included in this experiment. Pollen grain size and number were measured from five samples of each biological replicate using the Coulter counter Multisizer 4e. In this study, a mean of 10,948±2063 pollen grains were measured from each genotype (average 1121±208 pollen grains per plant) (Table 4). The three *Tazip4-B2* mutants showed a consistent and similar pollen profile comprising of two distinct peaks. The first peak represents pollen grains with grain size distribution similar to WT pollen and the second a group of pollen grains with smaller grain size (Fig. 6a). Accordingly, there were significant differences between the mean pollen grain size of each of the mutants *ph1b, ph1c* and CRISPR *Tazip4-B2* and their corresponding WTs (*P* < 0.01) (Table 4). More than 48% of pollen grains in the CRISPR *Tazip4-B2* hexaploid mutant samples were smaller in size (≤42 µm). A similar percentage of small pollen grains (47%) was found in the *ph1b* hexaploid mutant samples. However, small pollen grains (≤38 µm) were found in a lower percentage (34%) in the *ph1c* tetraploid mutant samples (Fig. 6b). The mean pollen number per anther ranged from 2317±333 to 3713±497 in the CRISPR *Tazip4-B2* and *ph1c* mutants respectively (Fig. 6c). However, no significant differences were observed between any of the *Tazip4-B2* mutants and their WT controls (Table 4). Detailed datasets of pollen size, pollen number per anther and percentage of small pollen grains for each *TaZIP4-B2* mutant and its respective wild type can be found in Table S7.

**Table 4.** Pollen number and pollen grain size for the three *Tazip4-B2* mutants and their corresponding wild types. Mean and median values with standard deviation are shown. Treatments with the same letter are not significantly different

| Genotype | Total number of measured pollen grains | N | Pollen grain size (μm) |  | Pollen number per anther |  | Small pollen grains (%) |  |
| --- | --- | --- | --- | --- | --- | --- | --- | --- |
|  |  |  | Mean | Median | Mean | Median | Mean | Median |
| *ph1b* | 10100 | 12 | 42.3 ± 1.3<sup>ab</sup> | 43.2 ± 1.7<sup>a</sup> | 2806 ± 426<sup>a</sup> | 2817 ± 430<sup>a</sup> | 47.2 ± 6.9<sup>a</sup> | 45.1 ± 8.3<sup>a</sup> |
| cv. Chinese Spring | 11072 | 12 | 47.4 ± 1.3<sup>c</sup> | 48.7 ± 1.5<sup>b</sup> | 3076 ± 370<sup>a</sup> | 3082 ± 364<sup>a</sup> | 15.0 ± 5.6<sup>b</sup> | 13.2 ± 2.5<sup>b</sup> |
| *ph1c* | 11139 | 10 | 40.9 ± 1.9<sup>b</sup> | 42.4 ± 2.6<sup>a</sup> | 3713 ± 497<sup>c</sup> | 3681 ± 593<sup>c</sup> | 34.3 ± 8.2<sup>c</sup> | 34.4 ± 8.3<sup>c</sup> |
| cv. Cappelli | 11840 | 10 | 43.7 ± 1.5<sup>a</sup> | 44.5 ± 1.5<sup>a</sup> | 3947 ± 270<sup>c</sup> | 3950 ± 248<sup>c</sup> | 11.1 ± 3.8<sup>b</sup> | 10.7 ± 4.0<sup>b</sup> |
| CRISPR *Tazip4-B2* | 13904 | 10 | 43.1 ± 2.1<sup>a</sup> | 43.4 ± 3<sup>a</sup> | 2317 ± 333<sup>b</sup> | 2354 ± 379<sup>ab</sup> | 48.5 ± 13.4<sup>a</sup> | 48.6 ± 13.4<sup>a</sup> |
| cv. Fielder | 11313 | 10 | 47.3 ± 1.0<sup>c</sup> | 49.1 ± 1.2<sup>b</sup> | 1886 ± 265<sup>b</sup> | 1875 ± 231<sup>b</sup> | 17.9 ± 3.9<sup>b</sup> | 19.1 ± 6.3<sup>b</sup> |

**Fig 6.**
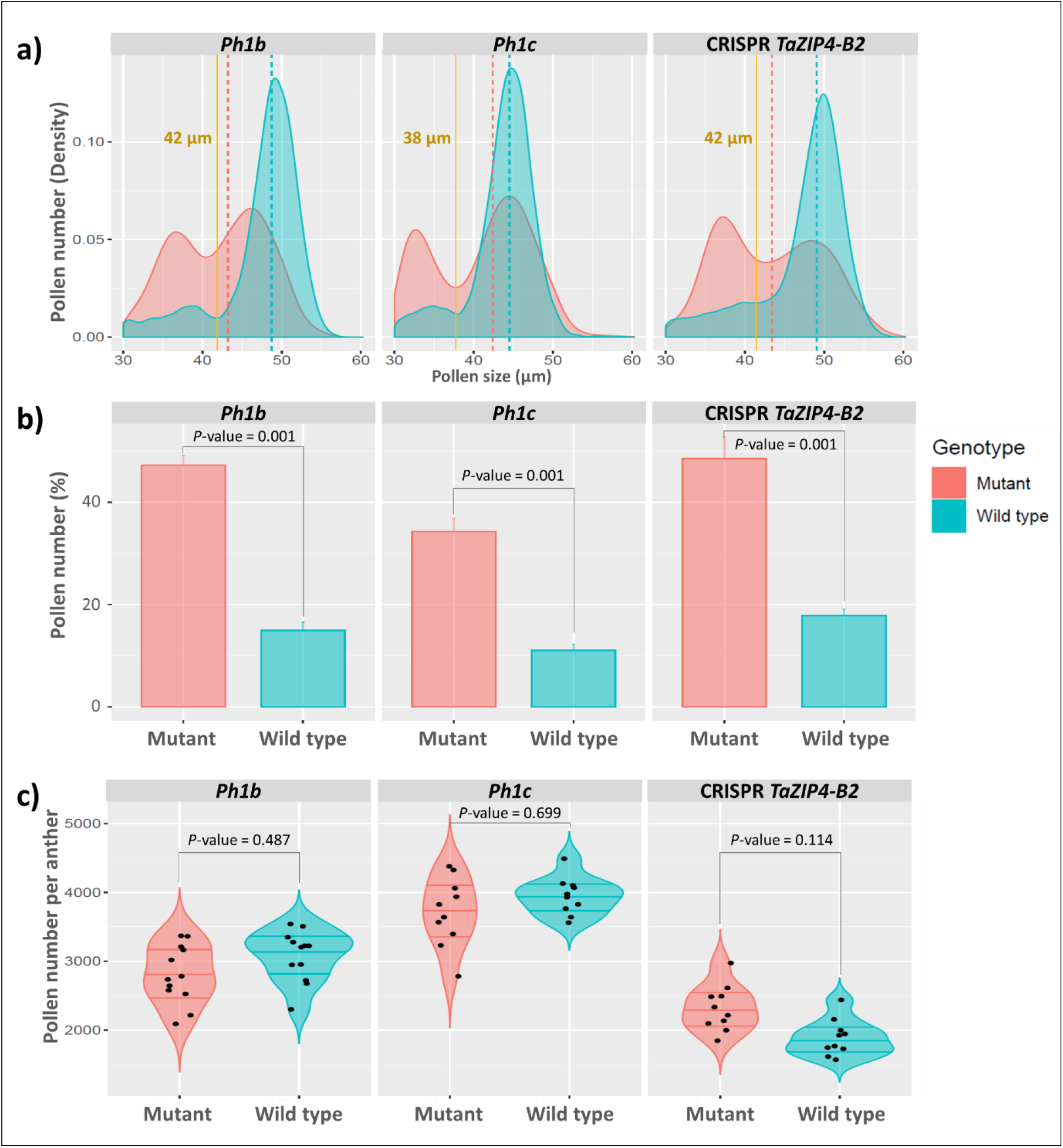
Pollen profiles of the three *Tazip4-B2* mutants. **a)** Density plot of the differential pollen size distribution data collected by coulter counter (Multiziser 4e) showing two distinguished peaks in all *Tazip4-B2* mutants comparing with their corresponding wild types. Dotted lines indicate the median pollen grain size for each genotype. Yellow lines indicate the borderline between normal and small pollen for each genotypes group. **b)** Percentages of the small pollen grains for each genotype (mutants and wild types). Pollen grain is considered small when it is ≤42 µm and ≤38 µm for hexaploid and tetraploid wheat pollen size, respectively. **c)** comparison of the number of pollen grains per anther between each *Tazip4-B2* mutant and its wild type. No significant difference in pollen number per anther was found between any of the mutants and its wild type.

Viability of pollen from the CRISPR *Tazip4-B2*, and *ph1b* hexaploid mutants, as well as the *ph1c* tetraploid mutant, was assessed using Alexander staining. More than 3000 pollen grains were scored for each genotype (from three biological replicates) after Alexander staining and image acquisition. Pollen coloured dark magenta after treatment with Alexander stain was considered viable, whereas light blue-green stained pollen was considered unviable (Fig. 7a). Analysis revealed similar percentages of unviable pollen grains in all *Tazip4-B2* mutants (Fig. 7c), with 28% in the CRISPR-*Tazip4-B2* mutant, 25.8% in the *ph1b* mutant and 22.8% in the *ph1c* mutant pollen being unviable (Table S8). In all cases, the level of unviable pollen grains in the mutants was significantly higher than that in the WTs (*P*<0.01), which did not exceed 3.3% on average. Developmental pollen stages were also assessed in the *TaZIP4-B2* mutants. Pollen grains from fully mature anthers from the Fielder mutant CRISPR *Tazip4-B2* and the WT Fielder were stained selectively for DNA using Feulgen stain. Results from the WT showed normal trinucleate pollen grains, whereas about half of the pollen grains in the mutant were immature and/or abnormal (Fig. 7b). Thus, the pollen profiling analysis revealed that around half the pollen from the CRISPR *Tazip4-B2* and *ph1b* hexaploid wheat mutants had similar pollen profiles, with around half the pollen grains being abnormally small. The Alexander and Feulgen staining methods provided further information revealing that the small pollen grains in the CRISPR *Tazip4-B2* and *ph1b* mutants are a mixture of both immature (unfunctional) and unviable pollen grains.

**Fig 7.**
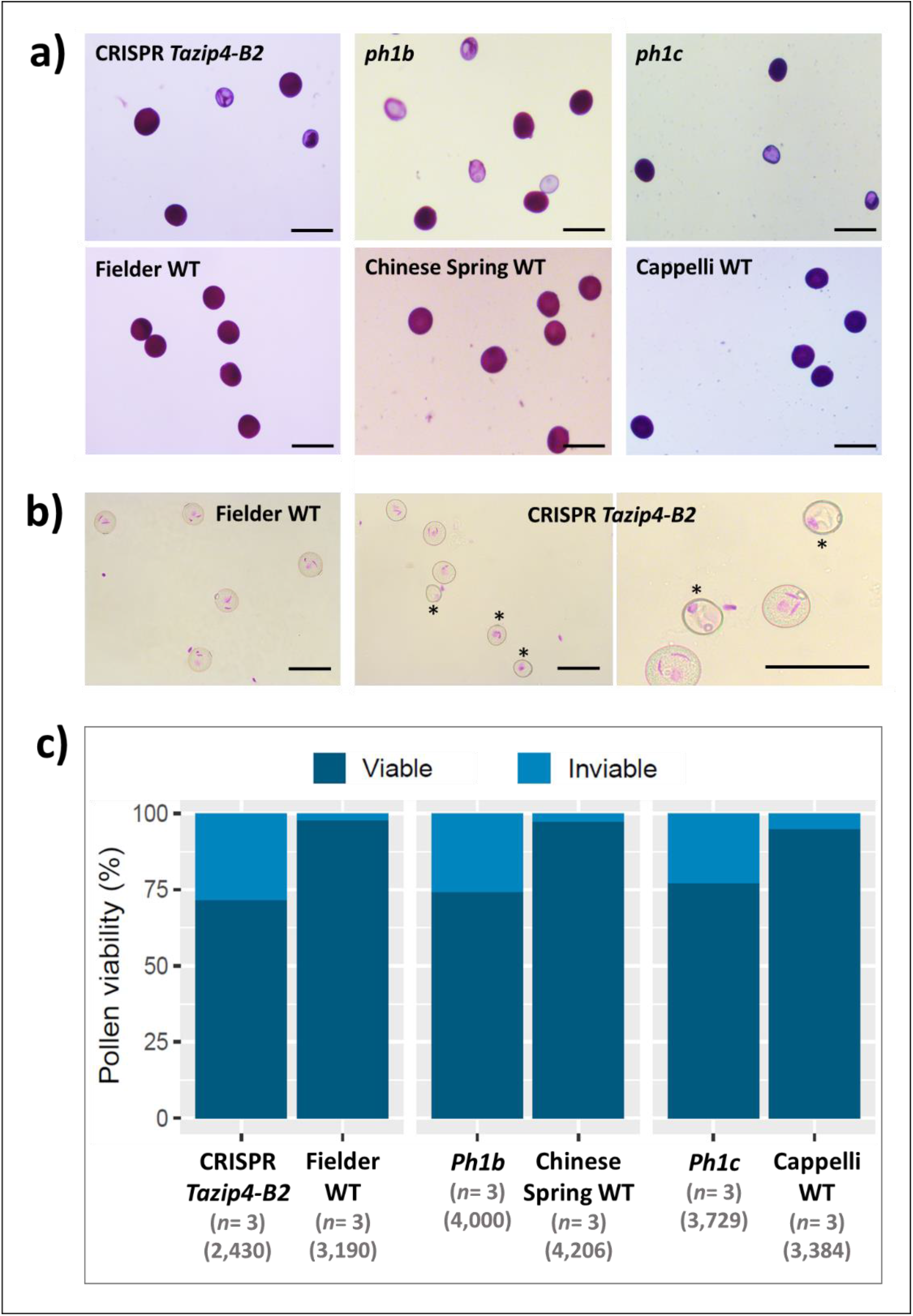
Pollen viability of the *Tazip4-B2* mutants. **a)**. Pollen with magenta colour after staining with Alexander stain was considered viable, whereas blue-green pollen was considered inviable. Bars equal 100 µm in length. **b)** Feulgen staining of pollens from anthers at anthesis in CRISPR *Tazip4-B2* mutant and its wild type (cv. Fielder) shows normal trinucleate pollen grains in the wild type, while almost half of the pollens were immature and/or abnormal in the mutant. Immature and abnormal pollen grains are indicated by an asterisk. Bars equal 100 µm in length. **c)** Percentages of viable and inviable pollens according to Alexander staining method for the three *Tazip4-B2* mutants and their wild types. *n* refers to the number of biological replicates. The numbers between brackets refer to the total number of scored pollen grains for each genotype.

## Discussion

Most polyploid literature highlights the requirement for a new polyploid to ensure the production of balanced gametes and hence fertility, through both meiotic and genomic adaptations (Comai, 2005; Otto, 2007; Pelé *et al*., 2018; Feliner *et al*., 2020). However, few meiotic adaptions have been characterised, which is surprising given their suggested importance to the preservation of overall polyploid fertility. Recent studies have implicated at least eight different autotetrapolyploid Arabidopsis meiotic genes in the meiotic stabilisation process (Yant *et al*., 2013), with specific alleles in one of these genes, *ASY3*, being highlighted in two further studies (Morgan *et al*., 2020; Seear *et al*., 2020). Allopolyploid Brassica studies have also reported a number of loci exhibiting natural variation in their ability to affect homoeologous pairing and crossover (Jenczewski *et al*., 2003; Liu *et al*., 2006; Higgins et al., 2020). Thus, these studies assessed the effects of natural gene variation on the meiotic process.

In the present study, we have assessed the stabilising effects of the meiotic gene *TaZIP4-B2*, which arose on chromosome 5B during wheat polyploidisation through duplication from 3B (Rey *et al*., 2017; Rey *et al*., 2018a; International Wheat Genome Sequencing Consortium, 2018). Results indicate that the deletion of the duplicated *TaZIP4-B2* copy (through CRISPR deletion of *Tazip4-B2*) results in 56% of meiocytes exhibiting meiotic abnormalities at metaphase I (Fig. 2; Fig. 3a, b); chromosome mis-segregation at anaphase I (Fig. 3c, d, e, f); 50% of tetrads possessing micronuclei (Fig. 3g, h, i, j); and finally, 48% of pollen grains being small (a mixture of immature and unviable) (Fig. 6; Fig. 7). A similar level of disruption is also observed in a hexaploid mutant (*ph1b*) carrying a 59.3Mb deletion encompassing *TaZIP4-B2*, with 56% of meiocytes exhibiting meiotic abnormalities (Fig. 2) and 47% of pollen grains being small (Fig. 6). Results suggest a direct correlation between meiotic abnormalities observed at metaphase I and pollen fertility. Importantly, there was also up to a 44% reduction in grain set in the CRISPR *Tazip4-B2* mutant (43% reduction in the *ph1b* mutant) (Fig. 4a). A considerable part of this reduction in grain set is likely to be due to pollination with immature/unviable pollen (Fig. 4b), rather than being mainly due to disruption in female gametogenesis. Pollen deposition and pollen grain size can have an effect on pollen competition for the ovule (Cruden & Miller-Ward, 1981; Németh & Smith-Huerta, 2003). However, it is still unclear how, within the 50:50 mixture of WT and immature/unviable pollen, WT pollen does not compete more effectively during pollination.

Development of *in situ* approaches are required to study the effect of *Tazip4-B2* on the female meiotic and post-meiotic stages. It will be particularly important to study the tetrad stage where only one megaspore survives (Morrison, 1953), to identify any preferential abortion of megaspores with unbalanced chromosome numbers resulting from disruption of meiotic pairing and crossover. However, whatever the importance of *Tazip4-B2* for female gametogenesis, the presence of *TaZIP4-B2* is still required to ensure nearly half the grain set in hexaploid wheat. This confirms the great importance and impact of the *ZIP4* duplication event on the fertility of this major global polyploid crop.

ZIP4 is a meiotic protein shown to be required for 85% of homologous crossovers during meiosis in Arabidopsis (Chelysheva *et al*., 2007) and rice (Shen *et al*., 2012). In polyploid wheat, the presence of *TaZIP4-B2* promotes homologous pairing, synapsis and crossover, and suppresses homoeologous crossover (Fig. 2) (Rey *et al*., 2018a). The deletion of *TaZIP4-B2* reduces homologous crossover (Rey *et al*., 2018a), contributing to an increase in meiotic abnormalities at metaphase I. The fact that the presence of *TaZIP4-B2* increases homologous crossover, suggests that the *ZIP4* effect on homologous crossover may be dosage dependent. This contrasts with the effect of other meiotic genes analysed in polyploid Brassica and wheat, where the loss of such genes does not reduce homologous crossover, but homologous crossover is only affected when all copies are deleted (Gonzalo *et al*., 2019; Desjardins *et al*., 2020). Thus, *ZIP4* was an effective target for divergence on polyploidisation, as its homologous crossover activity appears to be dosage dependent.

Although, *ZIP4* studies in Arabidopsis and rice have not shown a role for *ZIP4* in pairing and/or synapsis in these species, *ZIP4* is required for pairing and synapsis as well as homologous crossover in Sordaria (Dubois *et al*., 2019) and budding yeast (Tsubouchi *et al*., 2006). However, no *ZIP4* study in any other species has shown that it suppresses homoeologous crossover. This raises the question of how the duplicated *TaZIP4-B2* copy suppresses homoeologous crossover in wheat, and how it promotes homologous pairing, synapsis and crossover, preserving pollen viability and grain set. The early and 3-fold increased expression of *TaZIP4-B2* compared to the group 3 *ZIP4s*, is also likely to ensure that it competes with them for loading onto meiotic chromosomes (Rey *et al*., 2017). The present study reveals that up to half of the wheat ZIP4 protein is composed of TPRs (Fig. 1b, c). The presence of TPRs in other proteins has been shown to enable these proteins to form alpha solenoid helix structures (Blatch & Lassle, 1999; D’Andrea & Regan, 2003).

Previous studies have suggested that the *ph1b* deletion effect on homoeologous crossover in wheat is linked to the improved ability of the meiotic crossover protein MLH1 to process crossovers (Martín *et al*., 2014), while the *ph1b* deletion effect on chromosome pairing in wheat itself reported by Roberts *et al*., (1999) is linked to the chromosome axis protein, ASY1 (Boden *et al*., 2009). Recent studies in budding yeast have revealed that ZIP4 is connected to MLH1 through the binding of MER3, to ASY1 through the binding of another chromosome axis protein ASY3, and to synapsis proteins through ZIP2 (Pyatnitskaya *et al*., 2019). So, although *ZIP4* has not previously been shown to regulate homoeologous crossover in any species, or chromosome pairing and synapsis in plants, its interactions with axis and crossover proteins may provide a basis for these effects in wheat. Thus, the simplest explanation for the ability of *TaZIP4-B2* to promote homologous pairing and suppress homoeologous crossover, is that they result from a reduction in the normal functions of group 3 *ZIP4*s, as a consequence of the TPR divergence within *TaZIP4-B2* from that within *TaZIP4-B1* (Fig. 1b, c). The wheat group 3 *ZIP4*s are likely to process 85% of homologous crossovers as in other species (Chelysheva *et al*., 2007; Shen *et al*., 2012). They are also likely to process homoeologous crossover activity, given the level of crossover observed in wheat haploids lacking *TaZIP4-B2* (Jauhar *et al*., 1999). In contrast, although the diverged *TaZIP4-B2* copy has some homologous crossover activity, it does not possess any homoeologous crossover activity (Rey *et al*., 2018a). Sordaria studies reveal that the initial chromosome interactions involve ZIP4 foci on homologous chromosomes (Dubois *et al*., 2019). Thus, if wheat group 3 *ZIP4*s can process homologous and homoeologous crossovers, it is likely that foci of these ZIP4s can form stable interactions between both homologues and homoeologues. Again, the diverged *TaZIP4-B2* now only promotes homologous pairing or stable homologous interactions (Martín *et al*., 2018; Rey *et al*., 2018a).

Given the importance of the TaZIP4-B2 function for preserving grain number in wheat, future studies will need to confirm that the phenotype of *TaZIP4-B2* results from a reduction in the function activities possessed by the group 3 *ZIP4s*. Such studies will also need to confirm whether different *TaZIP4 B2* alleles exhibit variable phenotypes sensitive to temperature change. Natural variation in the meiotic phenotypes has been reported for some meiotic genes in other polyploids (Jenczewski *et al*., 2003; Liu *et al*., 2006; Yant *et al*., 2013; Morgan *et al*., 2020; Seear et al., 2020; Higgins *et al*.,2020).

The new approach for analysing pollen presented in this study can be used in future *TaZIP4-B2* studies to screen landrace diversity mapping populations (Wingen *et al*., 2017), carrying different *TaZIP4-B2* alleles for variable phenotypes, with variable sensitivity to temperature. This approach is high throughput and sensitive, with the capability to screen 1000s of pollen grains rapidly. Thus, the approach can be used for forward and reverse meiotic genetic screenings. In the present study, the technique was used to analyse pollen derived from both tetraploid and hexaploid meiotic mutants, revealing the presence of small pollen (Fig. 6a, b). The recent availability of multiple sequenced wheat genomes has allowed the initial identification of haplotype blocks (Brinton *et al*., 2020), revealing different *TaZIP4-B2* haplotypes. This information, combined with the availability of landrace diversity mapping populations (Wingen *et al*., 2017) and the pollen technique, can be used to rapidly identify any potential natural phenotype variation correlating with a specific *TaZIP4-B2* haplotype, as well as to explore the stability of such phenotypes under variable temperatures. This will be important for studies exploring the effects of temperature increases on wheat yields within the context of global climate change.

## Acknowledgements

This work was supported by the UKRI-Biological and Biotechnology Research Council (BBSRC) through a grant as part of the ‘Designing Future Wheat’ (DFW) Institute Strategic Programme (BB/ P016855/1) and Response Mode Grant (BB/R0077233/1).

## Author Contributions

AKA developed the pollen analysis and applied it to study the *Tazip4-B2* mutants. AKA undertook the grain set experiment; emasculation/pollination experiment, and their analysis, producing the figures and tables for all this data. AM carried out the cytological and immunolocalisation experiments and produced the immunolocalisation figure. GM carried out the TaZIP4 protein analysis, and AKA the sequence alignments producing the resulting figure. GM provided thoughts and guidance and revised and edited the manuscript produced by AKA and AM.

