## Supplementary material for "A duplicated copy of the meiotic gene *ZIP4* preserves up to 50% pollen viability and grain number in polyploid wheat": Fig. S1

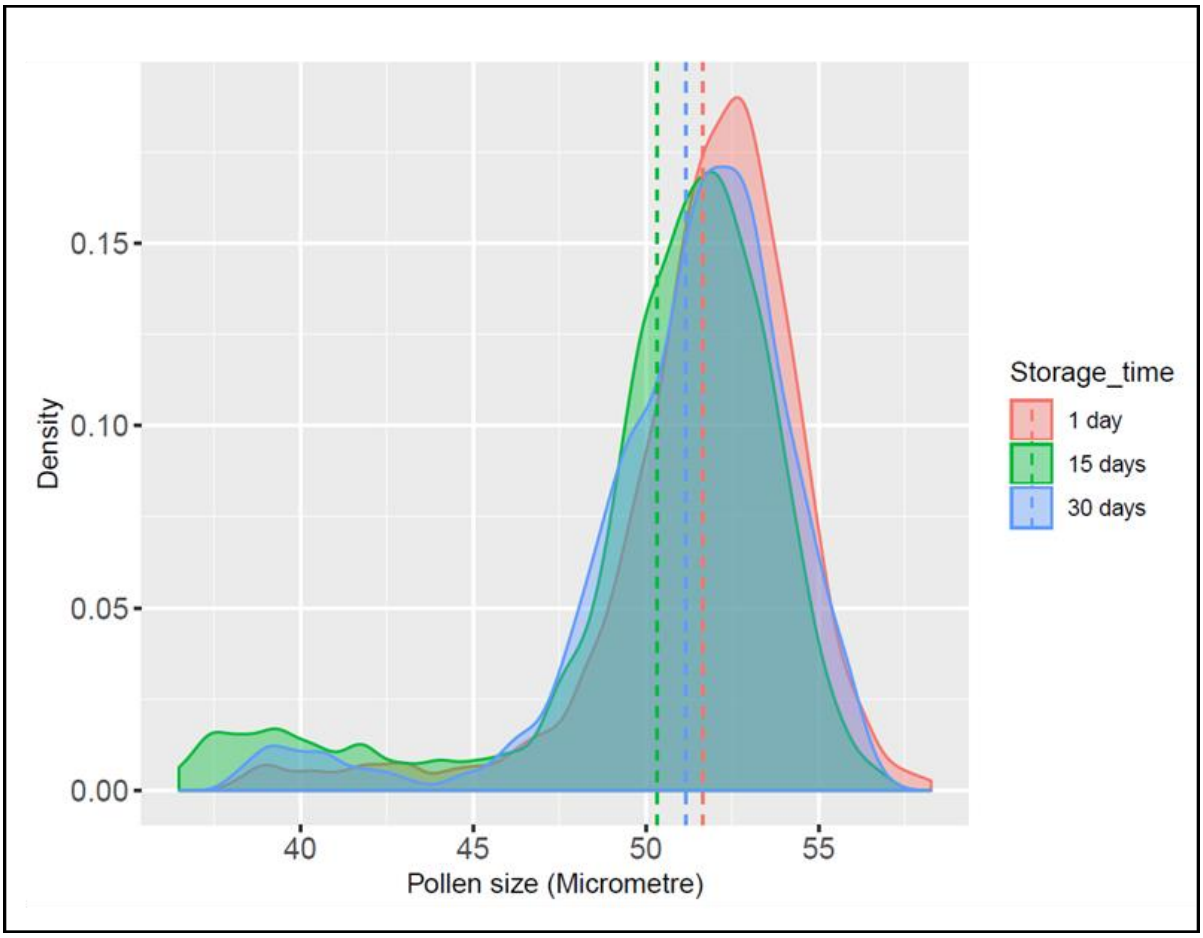


**Fig. S1. Comparing pollen profiles of same pollen samples after different storage times in 70% ethanol.** Five plants from hexaploid wheat (cv. Paragon) were used in this experiment. Three pollen samples (of three anthers from the same floret) were taken from the middle portion of the first spike of each plant. The three samples were mixed and homogenised and then divided to three equal parts. The first part analysed using the coulter counter (Multiziser 4e) after one day storage in 70% ethanol, while the second and third parts were analysed after 15 days and 30 days, respectively. The results showed that there is no significant effect of the storage time on the pollen size distribution pattern (pollen profile). Dotted lines indicate the mean pollen grain size for each treatment.
